# Spontaneous emergence of context-dependent statistical learning in humans and neural networks

**DOI:** 10.64898/2026.03.17.712206

**Authors:** Fleming C. Peck, Hongjing Lu, Jesse Rissman

## Abstract

Humans readily extract statistical regularities from experience, yet natural environments require flexible adaptation when associative structures shift across changing contexts, often without warning. Across two experiments, we show that humans can incidentally learn overlapping and conflicting visual associations even when contexts dynamically alternate and remain unsignaled or only minimally cued. To probe the computational mechanisms supporting this adaptive capacity, we trained recurrent neural networks with gated recurrent units on the same statistical learning task without providing any explicit context information. These models spontaneously developed distributed internal representations that robustly separated conflicting associations and supported rapid adaptation to latent context shifts. Critically, we show that these distributed representations, strongly shaped by the model’s initial weight configuration, played a key role in preventing catastrophic interference between contexts. Together, these behavioral and computational results significantly advance our understanding of how humans and artificial systems can successfully learn and flexibly retrieve context-dependent associations under challenging conditions.

## Introduction

Many everyday experiences unfold in structured, predictable ways, with events that recur over time in stable patterns. Internalizing these regularities allows anticipation of future occurrences, facilitating efficient information gathering, decision-making, and behavioral adaptation. It follows that the human brain is fundamentally oriented toward predicting the upcoming future based on recent events.^1–3^ This predictive ability helps conserve cognitive resources by reducing the need for continuous, effortful learning once patterns have been identified.^4^ However, the world is rarely static: associations often vary across contexts.^5^ To support adaptive behavior, the brain is thought to engage in context-dependent learning of these regularities and associations for flexible predictions as environmental conditions shift.^6^ For example, navigating a daily commute relies on learning the timing and location of traffic congestion, and expectations for social interaction may differ when a friend is encountered at work versus at a party. In both cases, prior experience supports the formation of context-bound predictions that guide perception and behavior.

Humans have an innate ability for statistical learning, allowing them to spontaneously discover regularities and associations. This process extracts spatial and temporal regularities from sensory input through passive exposure, without explicit instruction or external rewards.^7^ Statistical learning is proposed to support a wide range of cognitive functions, including language acquisition, visual perception, object recognition, and social cognition.^9,10^ Empirical studies demonstrate that individuals can detect regular patterns in continuous streams of stimuli across visual,^11^ auditory,^12^ and tactile^13^ modalities, in the absence of explicit transition cues and instructions.

Despite the rich literature on statistical learning, most research has focused on simple, highly reliable associations, such as detecting short sequences of objects or sounds. However, in natural environments, context plays a critical and often unobserved role in shaping how associations are formed. In animal learning research, for example, association-based behavior is known to be highly context-specific: extinguished fear responses return when animals are tested outside the extinction setting.^14^ Moreover, following extinction or reversal learning, animals reacquire original contingencies more rapidly than during initial learning,^15,16^ suggesting that prior contingencies are retained as latent knowledge in memory rather than being overwritten by new learning. Cognitive control processes are thought to underpin the behavioral flexibility afforded by suppressing previously useful but no longer relevant responses, allowing learners to pivot between contexts and contingencies as the environment demands.^17^ Notably, most studies in animal learning literature involve explicit reinforcement (e.g., reward or punishment), whereas statistical learning occurs incidentally without feedback, instruction, or overt motivation.

Although a few studies have explored the statistical learning of regularities that depend on a latent context or environment (e.g., ^18–22^), it remains unclear whether individuals can incidentally learn and retrieve context-dependent temporal associations without explicit perceptual context cues, reinforcement, or instruction. Analogous mechanisms have been proposed in sensorimotor learning, an instance of implicit learning where the brain is thought to infer context shifts and partition experience into distinct memories.^23^ Here, we test whether people can acquire two distinct sets of temporal associations instantiated with an overlapping pool of visual objects, where most associations are in direct conflict between contexts. For example, in Context A, Object X is followed by Object Y, whereas in Context B, the same Object X is followed by Object Z. Successful learning requires participants to flexibly update their expectations according to the active context inferred from recent sequence history. We examine how well human learners can discover these context-dependent associations without any external context cue – where context is embedded only in the pattern of transitions – using both offline testing and online learning measures.

To explore how these context-dependent representations might emerge from experience, we trained neural network models on the same behavioral task. We then identified the model that best matched human performance across the experimental conditions and analyzed its hidden-layer activations to generate testable hypotheses about analogous representations in the human brain. Deep neural networks have proven effective at capturing lower-level sensory processing,^24^ and recent perspectives advocate for extending these approaches to the study of higher-order cognition, including the representation of abstract knowledge.^25^ However, a common limitation of these modeling efforts is that these networks are typically trained on far more data than human learners (see ^26^ for a review), limiting the validity of direct comparisons. Additionally, prior modeling work frequently incorporates strong inductive biases that render context artificially explicit, either by feeding an unambiguous context signal into the input^22,27^ or by augmenting network architecture with designated units or computation modules.^28,29^ These modifications, while effective, constrain opportunities to observe how context discovery might emerge spontaneously. Inspired by Elman’s finding that simple recurrent neural networks can capture both short- and long-range dependencies^30^ and echoing recent calls to avoid hard-wiring solutions in cognitive modeling,^31^ we used minimally structured architectures that omitted context signaling and specialized inference modules. This design allowed us to examine how networks discover and represent latent task structure through sequence exposure alone. Finally, given evidence that weight initialization scale can influence learning trajectories in neural networks^27,32,33^, we systematically varied the initial weight magnitudes of the networks to assess how this factor affects their ability to learn and distinguish context-dependent associations.

The goal of the neural network modeling is to generate hypotheses about how the brain might represent context in statistical learning. The hippocampus represents two dominant neural representation strategies to support memory of individual experiences and to extract regularities across experiences^34,35^: sparse and distributed coding. Sparse codes, observed in the dentate gyrus and CA2/3 subregions, involve highly selective activation of a small subset of units in response to a given unit.^36^ Distributed representations, observed in the CA1 subregion, encode inputs across overlapping patterns of activity spanning the neural population.^37^ We specifically seek to find evidence of each of these strategies in the hidden layer activations of neural networks that successfully represent context-dependent associations.

Overall, this study aims to advance our understanding of context-dependent statistical learning by examining whether humans can learn and retrieve multiple conflicting statistical structures within highly overlapping stimulus sets. By manipulating the presence of visual contextual cues, we assess whether explicit signals of context shifts facilitate learning and whether individuals can still learn context-dependent associations in their absence. In parallel, we use neural network models trained on the same task to test whether artificial systems can account for human-like learning dynamics, offering insight into the computational mechanisms that may support flexible, context-sensitive learning in the brain.

## Results

Participants performed a context-dependent statistical learning task in which they viewed a continuous stream of 1,600 object images (Fig. 1A). Their only task was to indicate whether an “×” or “+” was embedded on each object (Fig. 1C), a perceptual judgment designed to maintain attention and allow tracking of online learning via reaction times (RTs). Unbeknownst to participants, the image stream was structured into object pairs specific to one of two distinct contexts. Although they were told that parts of the sequence might become familiar over time, they received no information about the underlying structure or the existence of multiple contexts. Each context defined a unique set of temporal associations between a largely overlapping object set, such that the probability of one object following another depended on the active context (Fig. 1B).

**Figure 1.**
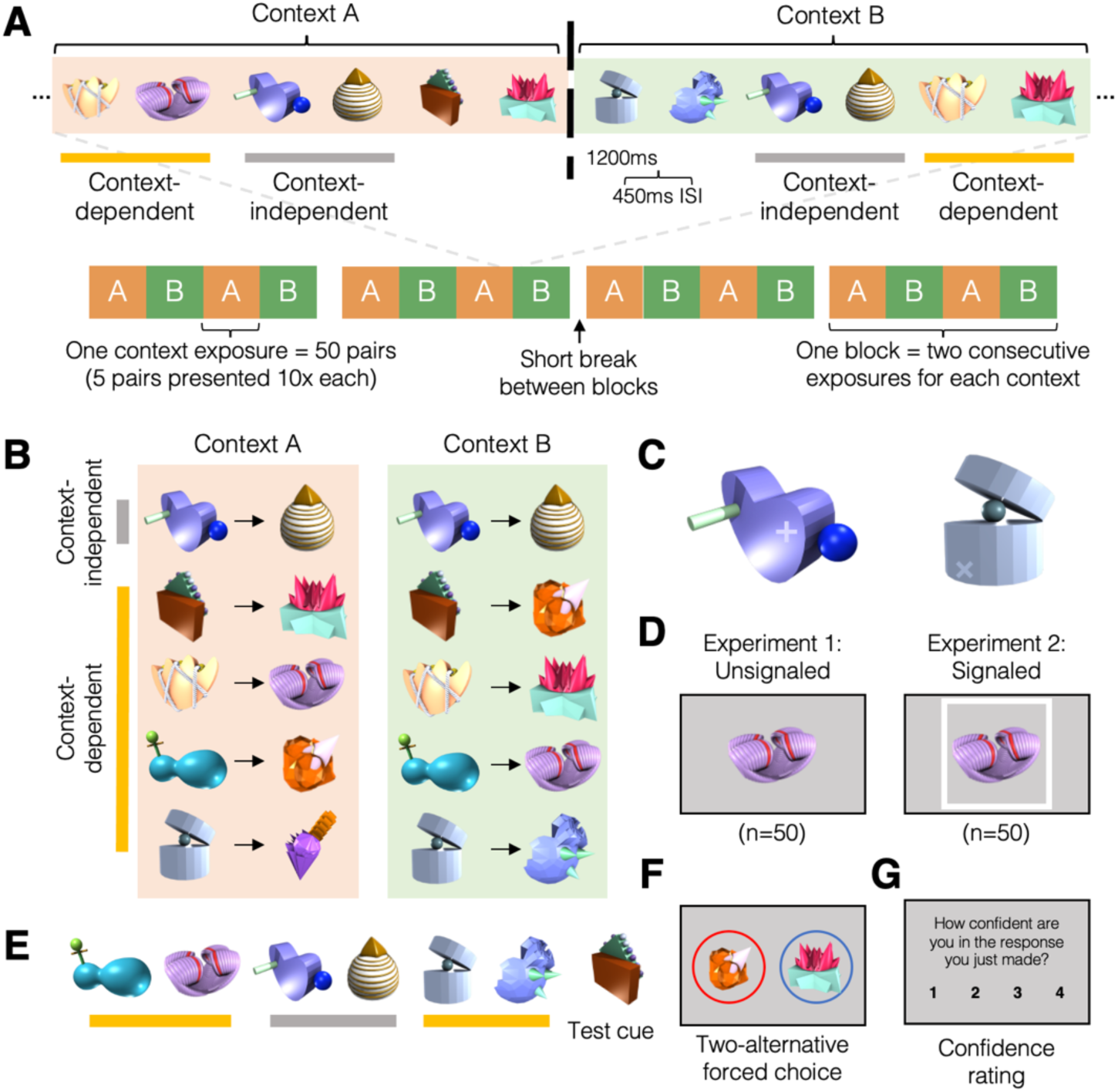
Experimental overview. *(A)* Visualization of the learning phase. Participants viewed a uniformly paced sequence of objects separated by brief fixation periods. Each object appeared for 1200ms with a 450ms interstimulus interval. The sequence was organized according to the temporal pair structure dictated by one of two contexts (Context A and Context B), which switched every 50 pairs. The orange and green backdrops are shown for illustrative purposes only. Participants performed four blocks of 200 pairs each, separated by short breaks. *(B)* Sample object assignments to context pair structures comprising 11 unique objects. The context-independent pair is the same for both contexts as shown in the first row, three of the context-dependent pairs consist of the same object set with pair assignment of the second pair position different for each context as shown in rows 2-4, and one context-dependent pair consists of a context-specific object in the second pair position as shown in the last row. *(C)* Example of object embedded with “+” or “×”. Participants were tasked with making a button-press response to indicate which symbol each object contained; object-symbol mapping was held constant throughout the experiment. *(D)* Differentiation of the two experiments: In Expt. 1 (Unsignaled), no context cues were shown and thus context switches were entirely latent (left); in Expt. 2 (Signaled), context was indicated with a white or black border around the object (right). *(E)* 2AFC test procedure: Example of 6-item (3-pair) sequence leading up to the test cue of a 2AFC trial. *(F)* Immediately following the test cue, participants chose which of two candidate objects comes next in the sequence. This example is a direct-conflict context-dependent trial, in which the lure corresponds to the object paired with the test cue in the other context. **(G)** After each choice, participants made a confidence rating.

Following the learning phase, participants completed a two-alternative forced choice (2AFC) test. Because objects appeared in both contexts, the correct association on a given trial depends on the active context. Accordingly, each test trial began with a six-object sequence composed of three object pairs from a single, consistent context, followed by the first item of a test pair (Fig. 1E). Participants were then tasked with choosing which of two objects should come next (Fig 1F). Context-independent trials assessed knowledge of the context-independent pair. Context-dependent trials consisted of two types: in direct-conflict trials, the lure was the object paired with the test cue in the other context; in indirect-conflict trials, the lure was an object not paired with the test cue in either context. After each choice, participants rated their confidence on a 1-4 scale (Fig. 1G).

In Experiment 1 (Unsignaled), n = 50 participants completed the task without any explicit perceptual context cue. In Experiment 2 (Signaled), a separate group of n = 50 participants completed the same task but with a visual context cue: a colored border (white or black) surrounding each object, corresponding with active context (Fig. 1D); this border was present during both the learning phase and the 2AFC test. Accuracy across the learning phase for the perceptual task was 92.2% for Expt. 1 and 92.7% for Expt. 2, indicating that participants attended to the stimuli during learning.

### Behavioral evidence of context-dependent statistical learning

We observed evidence of context-dependent statistical learning with significant 2AFC performance for both contexts (one-sample *t*-tests, Holm-Bonferroni corrected for three tests, all *p* < 0.001) (Fig. 2). When considering direct- and indirect-conflict 2AFC trials separately, we found above-chance accuracy for both trial types (all one-sample *t*-tests *p* < 0.05; *SI Appendix, Fig. S1*). A mixed-design ANOVA was conducted to examine the effects of experiment (Unsignaled vs. Signaled context, between-subjects) and context-dependence (context-independent vs. context-dependent trials, within-subjects) on 2AFC accuracy. There was a significant main effect of context-dependence (*F*(1, 98) = 17.52, *p* < 0.001), reflecting higher performance on context-independent than context-dependent trials. The main effect of experiment was not significant (*F*(1, 98) = 1.93, *p* = 0.17), nor was the interaction between experiment and context-dependence (*F*(1, 98) = 3.07, *p* = 0.08). We used Bayesian estimation to assess equivalence of context-dependent 2AFC accuracy between experiments. The posterior distribution of the mean difference was centered near zero (mean = 0.38%, 95% HDI [-4.1, 4.6]). Approximately 96.5% of the posterior mass fell within the predefined region of practical equivalence (ROPE) of [-5%, 5%], providing evidence that the two experiments yielded equivalent performance. This equivalence suggests that the border cue may have been too subtle to boost context-dependent learning or that explicit contextual cues are unnecessary to foster context-dependent learning beyond the contextual information that can be ascertained from recent sequence history in this paradigm. Additional analyses of confidence ratings and performance on remaining test tasks are reported in *SI Appendix, section S1*. These analyses reveal that most participants showed no explicit awareness of the temporal pair structure.

**Figure 2.**
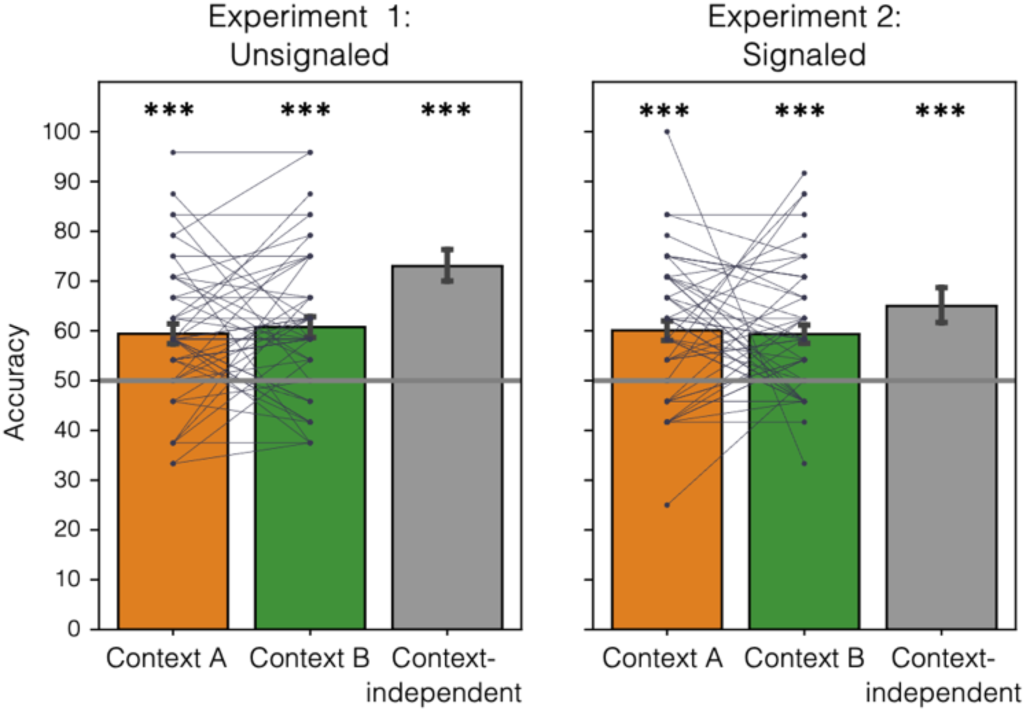
2AFC test performance. Bar height reflects group average 2AFC accuracy (% correct) for Context A questions (left bar, orange), Context B questions (middle bar, green), and context-independent questions (right bar, gray). Note that Context A and Context B correspond to the first and second contexts, respectively, used during the learning phase. Each dot reflects the accuracy for one participant with lines connecting a participant’s performance across the two contexts. Results plotted separately for Unsignaled and Signaled conditions on the left and right, respectively. Asterisks indicate significant deviation from chance performance (50%; horizontal line). ***p<0.001.

We also found evidence of an online learning effect using participants’ reaction times during the learning phase when they judged whether each object contained an “×” or “+” (Fig. 1C). Because these markers were consistently associated within corresponding objects across learning, faster responses could reflect memory-based predictions about the identity of upcoming objects consistent with rapid adaptation to temporal statistics in the sequence. We expect that over the course of learning, knowledge of the temporal pair structure would facilitate faster, anticipatory responses to the second item of each pair than the first item of each pair, which follows a random transition between pairs. Based on evidence that RTs improve throughout an experiment ^38^, we measure online learning as the second item RT subtracted from the first item RT, where a positive value indicates an anticipation effect, and a negative value reflects possible interference from context switches. Mean reaction times for each pair position are reported in *SI Appendix*, Table S1.

For context-independent pairs (Fig. 3; gray), we found a significant linear trend of RT differences across blocks in the Unsignaled experiment (*t*(49) = 2.70, *p* = 0.009), suggesting increasing anticipatory learning over time. However, no such trend was observed in the Signaled experiment (*t*(49) = 0.70, *p* = 0.49), where RT differences appeared to stabilize after the first block. For context-dependent pairs, both experiments showed a significant linear increase in RT difference across blocks (Unsignaled: *t*(49) = 4.36, *p* < 0.001; Signaled: *t*(49) = 2.59, *p* = 0.013). However, unlike the context-independent pairs, RTs for the predictable, item 2 objects in the Unsignaled experiment were initially slower than the first, unpredictable items (negative RT effect) and approached equivalence by the final block. This slowing earlier in the experiment may reflect interference from frequent context switches: participants had to suppress the prediction under the previously active context, which would be especially demanding during the early blocks of training. This effect is slightly ameliorated in the Signaled experiment, suggesting that participants may have been able to integrate the border contextual cue to facilitate online context-dependent learning. Despite this initial disadvantage for second item responses, the online learning measure increased over time, reaching its highest average in the final block. A mixed-design ANOVA on the RT difference score with experiment as a between-subjects factor and block as a within-subjects factor showed no main effect of experiment (*F*(1, 98) = 1.48, *p* = 0.23).

**Figure 3.**
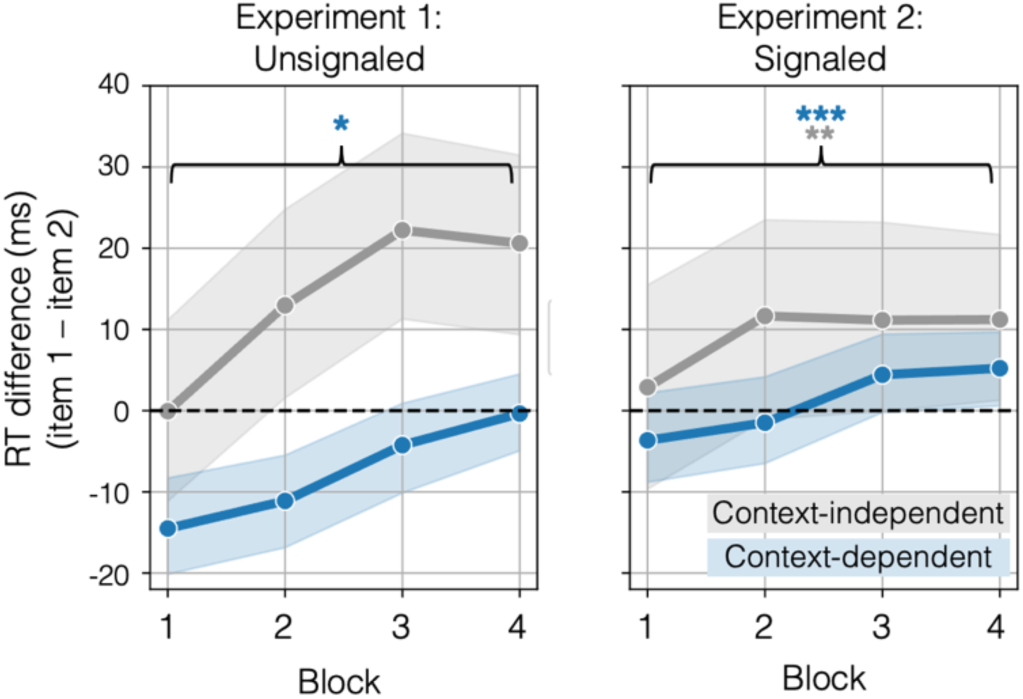
Reaction time differences reveal trajectory of online learning. Reaction time (RT) difference between responses to objects in the first (item 1) and second (item 2) pair position. A positive value on the y-axis shows anticipation effect plotted for each block during learning phase (x-axis). Average RT difference with standard error of the mean (shaded) for context-independent pairs in gray and for context-dependent pairs in blue. Linear trend significance indicated with same color scheme. Linear contrast significance indicated, ***p<0.001; **p<0.01; *p<0.05. The shaded areas indicate sampling error.

### Neural network weight initialization influences context-dependent learning

Having established that humans can spontaneously learn context-dependent associations from exposure alone, we next turned to a computational account of this behavior using artificial neural network models. Our goals were to test whether these models could similarly discover the task’s latent structure without context cues and, critically, to characterize the nature of the emergent representations that give rise to the context-dependent gating of associative predictions, examining how specific model parameters shape this capacity.

First, we determined that recurrent neural networks with gated recurrent units (GRUs) learned the task more effectively than other network architectures, including feedforward networks and recurrent networks without gated units (see model comparison details in *SI Appendix*, section S2). Next, we trained GRU models on the same amount of sequence exposure as human participants. Models featured a 150-node hidden layer and were trained to predict the next item in the sequence using one-hot encoded object representation for both inputs and outputs (Fig. 4A). Critically, models received no explicit context information, requiring them to discover the latent structure from sequence statistics to make accurate predictions. As with humans, learning was evaluated with a 2AFC test. Model weights were frozen after training, and each 2AFC trial presented the model with a series of seven objects (i.e. a three-pair sequence and a test cue).

**Figure 4.**
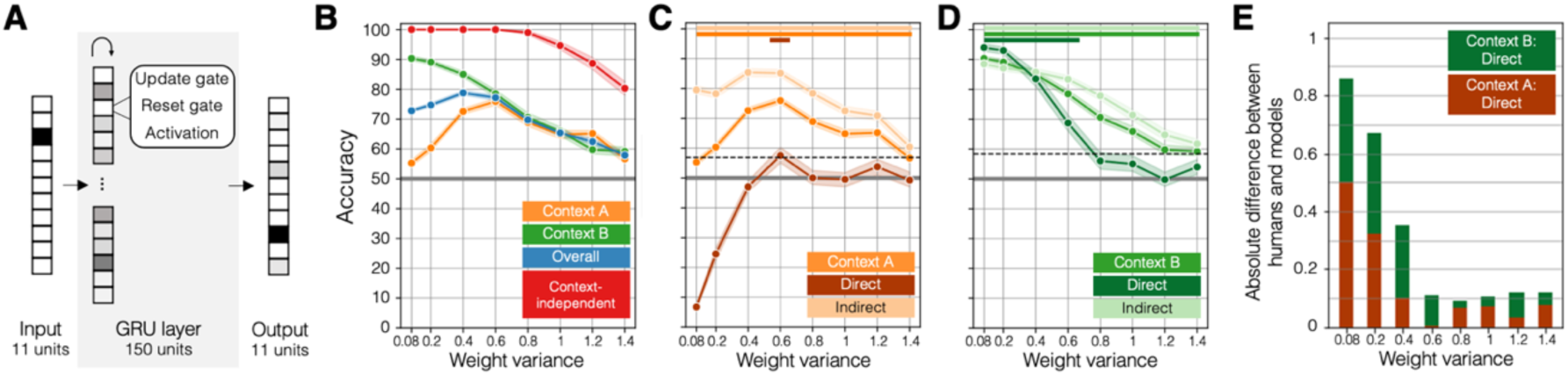
GRU model’s 2AFC performance by weight variance. *(A)* Visualization of neural network architecture comprised of 11 input units, a single GRU layer with 150 units, and 11 output units. *(B-D)* 2AFC accuracy (y-axis) on context-dependent test trials for GRU models with weights initialized with increasing variance along the x-axis color-coded by question category. *(B)* 2AFC performance on Context A (orange), Context B (green), overall context-dependent (blue) and context-independent (red). Chance performance (50%) indicated with gray horizontal line. *(C-D)* 2AFC performance on individual contexts, visualized for overall as well as direct-conflict (dark coloring) and indirect-conflict (light coloring) trial subsets. Significant one-sample *t*-tests from chance (Bonferroni-corrected for eight comparisons) indicated with horizontal lines at top of plot color-coded in the same way. Mean human performance on direct-conflict trials of each context indicated by the dashed horizontal black line. *(C)* Context A. *(D)* Context B. *(E)* Absolute difference direct-conflict 2AFC performance between human group average and model group average for each weight initialization configuration. Bar height reflects summed direct-conflict 2AFC performance absolute difference of Context A (dark orange) and Context B (dark green).

The model then “selected” the next object in the sequence between two options, with its choice determined by the object with the higher predicted probability.

We systematically varied the bounds of the uniform distribution used to initialize model weights to evaluate whether greater initial weight variance would accelerate convergence, motivated by prior findings that initialization in neural networks can strongly influence learning dynamics.^39,40^ Low-variance initialization is commonly used as the default in neural networks. However, it remains unclear whether this default choice affects a model’s capacity to learn latent structures in the data. To address this, we systematically varied the weight initialization variance across a wide range of values. For each weight variance initialization condition, we trained and tested 50 independent models and report the average performance.

Across weight initialization conditions, models with low to moderate initialized weight variance achieve perfect accuracy on context-independent trials, demonstrating their ability to learn stable, non-contextual associations (Fig. 4B). However, as variance of initial weights increases, performance steadily declines, highlighting how excessive initial weight variance introduces noise, disrupting the model’s ability to extract consistent patterns from the sequence.

The models’ learning of context-dependent associations – where context-specific conflicts must be resolved – reveals more complex dynamics. The relationship between initialized weight variance and context-dependent accuracy is non-monotonic, with the highest performance demonstrated by models with moderate weight initialization variance within the range of (0.4-0.6) (Fig. 4B). Low-variance models (0.08-0.2) demonstrate around 90% accuracy on Context B, the context which the model was most recently processing at the end of training before model weights were frozen, compared to below 60% accuracy for Context A, the context which was previously learned and conflicted with the more recently learned associations. Increasing initialized weight variance in the high-variance range (1.0-1.4) exhibits a steady decline in accuracy for both contexts, indicating that their representations may be too diffuse or unstable, preventing them from effectively differentiating between contexts. Notably, this non-monotonic relationship between 2AFC accuracy and initialized weight variance was preserved when models were trained using input representations that reflected the perceptual features of the objects (as opposed to one-hot vectors), derived from computer vision models such as AlexNet (*SI Appendix*, section S3). The same pattern held when the context-dependent pair that included a context-unique second item was excluded, such that all pairs comprised items that appeared in both contexts (*SI Appendix*, section S4). This result helps rule out the possibility that recent exposure to particular items served as a context cue, strengthening the interpretation that context is inferred from recent sequence history.

Breaking down performance into direct-conflict and indirect-conflict trials reveals notable differences in model learning. Direct-conflict trials (where the lure object is the correct answer for the other context) are the most diagnostic test of context-dependent learning as they place associations from different contexts in direct competition, making accurate performance dependent on the use of contextual information to disambiguate the correct response. Only models initialized with a weight variance of 0.6 achieved above-chance performance on Context A direct-conflict trials (*t*(49) = 2.81, *p* = 0.004), though this came at the expense of reduced though still significant accuracy on Context B direct-conflict trials (Fig. 4D). Low-weight models perform near floor on Context A direct-conflict trials, rendering their high overall context-dependent accuracy misleading as it reflects only strong performance on indirect-conflict Context A questions and mastery of Context B. In contrast, high-weight models show no advantage for Context B direct-conflict trials, with performance on direct-conflict questions around chance for both contexts.

To identify which weight initialization variance best approximated human-like behavior, model performance on Context A and Context B direct-conflict trials was compared to human data. Human accuracy was averaged across the Unsignaled and Signaled experiments as no significant difference was observed between them. Fig. 4E visualizes the absolute difference in 2AFC performance between models and humans on direct-conflict trials for both contexts (human performance indicated by the dashed black line in Fig. 4C, 4D). Initialized weight variance of 0.6 produced the clearest match to human performance, forming an elbow in the plot and achieving above-chance accuracy across all question sets.

### Distributed hidden layer representation strategy facilitates context-dependent learning

Having observed that models with initialized weight variance in the moderate range (such as 0.6) successfully learned context-dependent associations without any explicit context signal as input, we next examined how context information is encoded in the hidden layer activations of the models. Context encoding could manifest as either a sparse representation (carried by a few units) or a distributed representation (spread across many units). Therefore, we quantified two complementary properties of the activations: the extent to which context sensitivity was localized to a small subset of units (akin to individual “context cell” neurons that code for which context is currently active), and the degree to which the currently active context was expressed a as distinctive pattern of activity across many units. These analyses were conceptually motivated by prior work distinguishing sparse and distributed coding strategies in hippocampal and connectionist models.^35,41^ The analysis was conducted using the hidden-layer activations during the final block of training.

The *sparse representation index* measures the proportion of hidden layer units that do not show significant activation differences between contexts. A higher sparse representation index therefore indicates that fewer units selectively encode a specific context, while a larger proportion of units are context-insensitive (Fig. 5A). This measure of context sensitivity for each unit was calculated with a one-way ANOVA comparing activations during exposure to Context A versus Context B in the final quarter of training (block 4). Fewer nodes with significant context sensitivity indicate that limited number of hidden-layer units support the context-specific representations.

**Figure 5.**
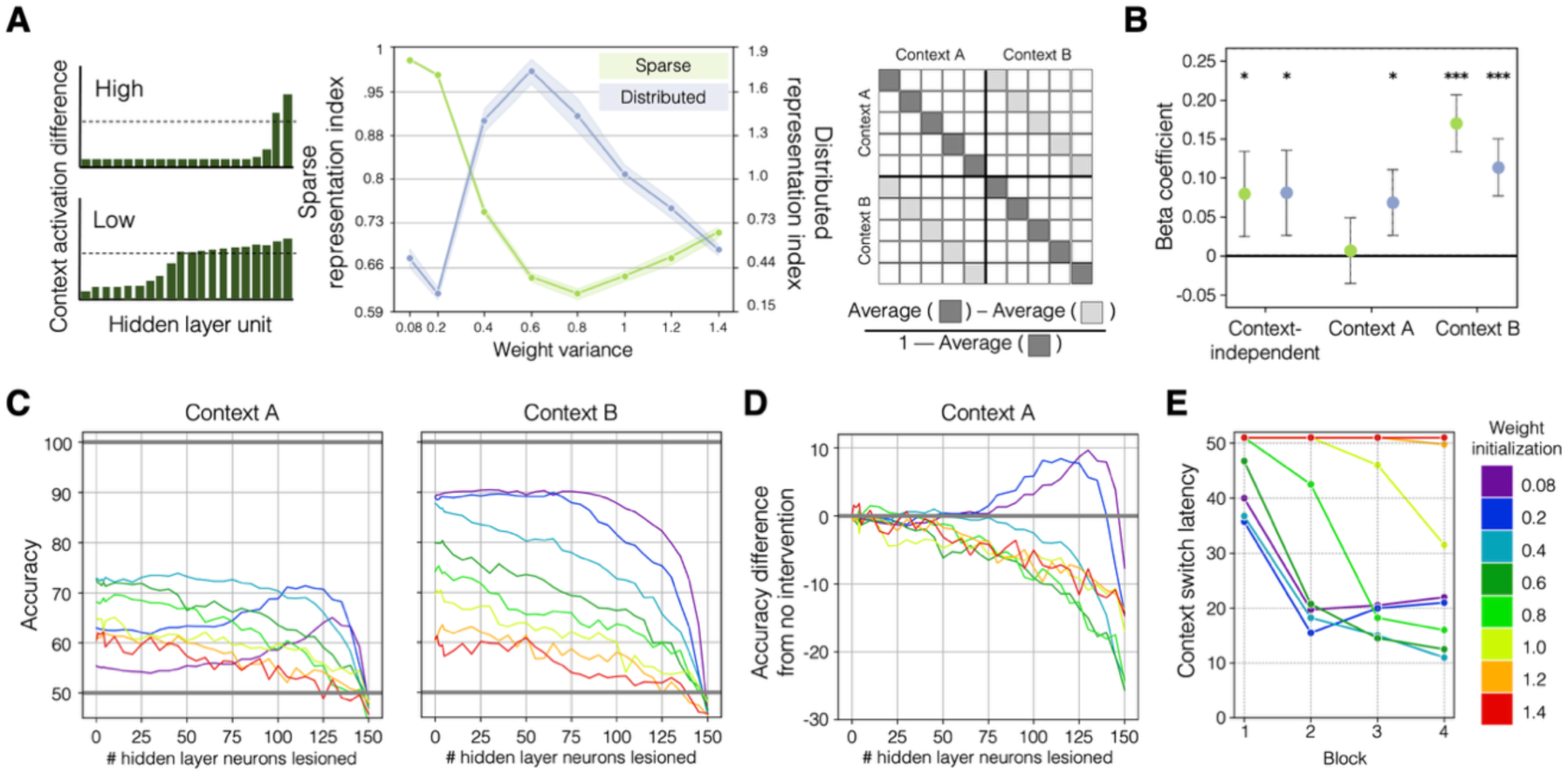
Neural network hidden layer task representation strategies. *(A)* Visualization of computation of sparse representation index (left) and distributed representation index (right) plotted for each weight variance configuration (middle; x-axis) in light green and blue, respectively. *(B)* Significance of beta coefficients (y-axis) for multivariate regression analyses using sparse (green) and distributed (blue) representation indices to predict each 2AFC question category (x-axis). ***p<0.001; **p<0.01; *p<0.05. *(C-D)*: Lesion analysis results. *(C)* 2AFC accuracy of Context A (left) and Context B (right) questions (y-axis) as an increasing number of hidden layer nodes are lesioned in descending rank order of context sensitivity (x-axis) for each weight initialization configuration (rainbow coloring). *(D)* 2AFC accuracy performance difference from no intervention for lesion analysis. *(E)* Context switch latency (y-axis) visualized across learning phase blocks (x-axis) for each weight initialization configuration (rainbow coloring). Failure to reflect a context switch is noted with a value of 51 (e.g., greater than duration of context exposure).

The *distributed representation index* was derived from a representational similarity analysis of hidden layer activations for the first item of each pair, the item that carries the context-dependent association. The index compares the geometric distance (dissimilarity) of these representations within a context versus across contexts, with normalization based on within-context consistency so that more stable representations are given greater influence (Fig. 5A, right plot). Higher values indicate that distinct context representations are distributed across the hidden layer.

We found that the low-variance models exhibit the sparsest representations, and the moderate-variance models exhibit the most distributed representations (Fig. 5A, middle plot). High-variance models do not show strong evidence of either representation strategy. The 0.6-initialized model, which was the only model to demonstrate significant direct-conflict 2AFC accuracy for both contexts (Fig. 5B-C), exhibited the strongest evidence for the distributed over the sparse representation index.

To understand how these representation strategies supported learning, we regressed 2AFC accuracy on the z-scored sparse and distributed representation indices (averaged across the 50 models initialized for each weight configuration; Fig. 5B). Including both predictors in the same model allows us to evaluate their unique contributions to 2AFC task performance. Both indices significantly predicted context-independent accuracy (Sparse: β = 0.080, *p* = 0.016; Distributed: β = 0.081, *p* = 0.015). For context-dependent learning, the sparse representation index predicted performance only for Context B (β = 0.17, *p* < 0.001) but not Context A (β = 0.007, *p* = 0.71), consistent with its prominence in the low-variance models that disproportionately learned Context B. In contrast, the distributed representation index predicted accuracy for *both* Context A (β = 0.069, *p* = 0.012) and Context B (β = 0.11, *p* < 0.001), reinforcing that this strategy more effectively supports context-dependent learning, where successful learning requires retaining knowledge of both contexts.

### Efficient context switching facilitates expression of context-dependent knowledge

We next carried out a lesion simulation analysis to understand how the moderate-variance models provide a better account of human behavior than the low-variance models, which is often used as the default initialization in neural networks. For each model, all 150 hidden layer units were ranked according to their context sensitivity index (the *F*-statistic of activity difference between Context A and Context B). We then progressively lesioned the most context-sensitive units by setting their activations to zero and re-evaluated 2AFC accuracy after each lesion step.

The moderate-variance models show a steady decline in both Context A and Context B accuracy as more nodes were lesioned (Fig. 5C). This result further indicates a distributed representation strategy where many units contribute uniquely to the representation of current context. In contrast, the low-variance models show evidence of a redundant coding strategy: accuracy, particularly for Context B, remains largely unchanged until around half of the hidden layer was lesioned (Fig. 5C). This delayed performance decline complements the earlier finding of a sparse representation strategy, in which very few nodes showed significant context sensitivity, suggesting that most units carried only shallow, overlapping context signals. Then, when Context B performance began to decline, Context A performance actually increased (Fig. 5D), with performance eventually reaching level comparable to the moderate-weight models (Fig. 5C). This result indicates that the Context A representations are present in the knowledge base of the network. However, the context knowledge is not accessible for the 2AFC testing task given the low-variance models show excellent Context B accuracy (the context on which it was more recently trained) but extremely poor Context A accuracy (the previous context) (Fig. 4A).

To explain the discrepancy in 2AFC accuracy in light of this evidence of Context A knowledge preserved in both low- and moderate-variance models, we examined how efficiently models adapted to context switches. We derived a context switch latency metric operationalized as the number of pairs the model processed after a context switch before perfectly predicting the paired associate of all remaining pairs in each 50-pair context exposure. We found that, by the final block of training, the moderate-variance models exhibited faster switch latencies whereas the low-variance models adapted more slowly (Fig. 5F). This inefficiency likely prevented the low-variance models from shifting away from their end-of-training state during the brief six-item exposure provided in each 2AFC trial, leaving them biased toward Context B despite evidence of retaining earlier Context A associations. Overall, this evidence indicates that the moderate-variance models achieve the best context-dependent learning because they retain associative knowledge across contexts and quickly adapt to context switches with support from a distributed representation of context across the hidden layer.

## Discussion

Given that statistical learning enables associative strengths to be incrementally updated over many exposures, it has been unclear whether it affords sufficient flexibility to adapt to changing associative contingencies in different contexts. The present results extend our understanding of when incidental learning of temporal regularities is possible via demonstration of context-dependent statistical learning under circumstances where contexts dynamically alternate, associations directly conflict, and no explicit instructions are provided. We found evidence for this in above-chance performance on the final 2AFC test as well as progressive RT speeding for predictable objects compared to unpredictable objects over the course of learning, which suggests that implicit learning mechanisms facilitated anticipatory behavior.^42^

Notably, explicitly signaling context with a colored border (Expt. 2) did not enhance context-dependent learning compared to when context was fully latent (Expt. 1). This may reflect greater influence of local temporal context (e.g., recent sequence history, which was available during both learning and retrieval) over environmental cues or disruption of implicit learning mechanisms by promoting a more deliberate strategy. Indeed, recent work suggests that states of reduced executive control, such as mind wandering, can enhance statistical learning relative to focused on-task states,^43^ implying that exogenously focusing attention via explicit cues may be counterproductive for this type of incidental learning. However, given that the border cue changed color only every few minutes and its relevance was not explicitly conveyed, it is also possible that this visual cueing of context may have been too subtle to provide a performance advantage; a more salient context signal might have produced different effects.

Prior efforts to demonstrate context-dependent statistical learning with auditory stimuli have been unable to find learning of both contexts unless participants were provided with explicit instructions or salient context cues.^18,19^ Our success in the visual perceptual domain supports accounts suggesting that statistical learning mechanisms may be modality-specific rather than fully domain-general,^44^ with visual statistical learning potentially more robust to context-based interference under implicit learning conditions. Siegelman et al.^20^ reported some evidence of context-dependent learning in the visual domain using associative structures built from an overlapping set of stimuli. However, their paradigm involved a single consecutive exposure to each of the two contexts, rather than repeated interleaved context switching, self-paced stimulus sequence exposure, and explicit instructions to look for patterns in which shapes tended to follow each other. Such design choices are different from the present study that focuses on shorter, fixed-duration stimulus presentations to minimize possibilities for strategic encoding and support the passive, implicit learning that is thought to characterize statistical learning.^45^

The present findings build on prior work on second-order conditional (SOC) sequence learning, which has demonstrated that learners can extract higher-order temporal dependencies in which predictability depends on combinations of preceding elements rather than simple pairwise transitions.^42,46^ Recent work further suggests that exposure to SOC structure can shape subsequent performance and subjective sensitivity to sequence regularities even when explicit knowledge is limited.^47^ Although the surface structure of these tasks differs from the present paradigm, both lines of work underscore how context-sensitive behavior can emerge from the integration of temporal regularities over experience, without requiring explicit contextual signals.

We used a neural network modeling approach to inform hypotheses of how the human brain might support such learning. These models were optimized to predict the next object in the sequence. While our human participants engaged in a cover task requiring simple ×/+ perceptual judgments, we assume that they were implicitly forming predictions about upcoming stimuli. Therefore, the models’ predictive framework captures a core computational goal that the human learners pursue implicitly: anticipating future input based on recent experiences.^1^

The GRU’s gating architecture may support its successful context-dependent learning by enabling the model to manage conflicting associations based on retaining relevant information while filtering out noise. This computational function parallels how the human brain manages interference between new and old memories.^48^ Although GRU models are not intended as models of biological mechanisms, the update and reset gates bear resemblance to the dynamic interplay between the hippocampus and neocortex that supports stability for long-term memory storage^34^ as well as to neuromodulatory systems where prediction error signals (i.e., dopamine release) prompt a reassessment of context and switch in behavioral strategy.^49^ Such mechanisms have been hypothesized to facilitate segmenting continuous experiences and recalibration of predictions,^50^ which may be relevant for context-dependent temporal associative learning and motivate hypotheses for future work examining parallels between biological systems and computational models.

Prior neural network modeling of context-dependent learning has imbued neural networks with specialized architecture to facilitate latent cause inference.^28,29^ or have explicitly provided unambiguous context information in model input.^22,27^ For example, Smith and colleagues^51^ effectively demonstrated that recurrent networks can track temporal structure across multiple timescales within explicitly signaled contexts in a statistical learning paradigm instantiated as games that share response choices. However, these studies bypass the question of how a sense of context might emerge organically from exposure alone to disambiguate overlapping task structure. Additionally, they introduce assumptions that are arguably biologically implausible, such as constant context monitoring and perfectly reliable context cues.^5^ Here, we more directly focus on latent context discovery by exploring how weight initialization affects learning dynamics. Since network weights are adjusted throughout training to minimize loss, their initial configuration acts as a key driver of convergence.^52^ Prior work suggests that higher initial weight magnitudes bias models toward “lazy” solutions, involving rapid solution convergence with unstructured representations, while smaller magnitudes support “rich” solutions that exhibit more structured learning albeit at a slower pace.^27,33,40^

Indeed, increasing the variance of the uniform distribution used to initialize model weights to a moderate range facilitated successful context-dependent 2AFC performance. This improvement was associated with a high-dimensional, distributed code in the hidden layer that was significantly associated with 2AFC trials of both contexts. This is consistent with studies suggesting that high dimensional codes afforded by mixed selectivity in prefrontal cortex neurons allow for more flexibility and rapid adaptation to new tasks.^41,53^ The successful distributed context coding strategy where identical model input is represented differently when processed in different contexts is consistent with reports of the hippocampus integrating contextual information into stimulus representations.^54,55^ Furthermore, the hippocampus supports the rapid learning of temporal associations.^37,56^ Taken together, these parallels suggest that the moderate-variance GRU models are capturing both higher-level contextual encoding and lower-level temporal associations, consistent with core functions of the hippocampus.

The variance of weight initialization may be interpreted as shaping the GRU’s inductive bias: the assumptions the model makes about the structure of the environment, particularly regarding the presence and separability of underlying contexts. Low initial weight variance appeared to bias the model towards rigid representations that emphasize on recently experienced associations and failed to recover earlier learned patterns following context shifts. On the other end, high initial weight variance produced overly flexible representations that failed to consolidate stable structure. Our analyses suggest an optimal intermediate range of initial weight magnitudes, where models were sufficiently flexible to distinguish between contexts yet structured enough to preserve associations within each context and avoid catastrophic interference. Accordingly, these effects are best understood as emergent inductive biases shaped by properties of training dynamics and initialization, which may provide insight into how learning systems come to represent and segregate latent contexts, whether biological or artificial. Future work could assess the extent to which similar learning dynamics arise across architectures and task demands and whether manipulating network hyperparameters, such as learning rate and number of hidden layers, consistently shape the balance plasticity and stability in context-dependent learning.

To better understand why the moderate variance models succeeded, it is informative to examine the limitations of the low variance models. These models performed poorly on Context A test trials, a pattern that might initially suggest catastrophic interference – that previously learned associations of Context A were overwritten by more recent Context B experience. However, the lesion analysis revealed that Context A knowledge remained in the networks but was not accessible until over half the hidden layer was removed. One likely explanation for this inaccessibility is the slower context switching in low variance models: compared to the moderate variance models, which successfully expressed knowledge of both contexts, the low variance models were slower to accommodate context switches. As a result, the brief context exposure sequences preceding each 2AFC decision may not have provided sufficient evidence to pull them out of their orientation towards Context B state at test, which remained simply because Context B was the last context encountered during training. This phenomenon parallels findings from the fear extinction literature, where extinguished fear responses can re-emerge in a different context, indicating that underlying knowledge is retained but not manifested in behavior when irrelevant to current setting.^14^

Another key limitation of the low variance models that emerged from the lesion results was a constraint on how knowledge was represented in the hidden layer units. Before Context A performance recovered, these models showed little to no change in 2AFC accuracy for either context until roughly half the hidden layer was lesioned, in contrast to the steady performance decline observed in moderate and high variance models. This suggests highly redundant coding within the hidden layer. Redundant neural coding is theorized to enhance robustness in noisy environments by duplicating information across neural populations^53,57,58^ – a potentially advantageous feature for the present task, where many associations directly conflict and half of the training samples are unreliable (e.g., between-pair transitions). Such redundancy could plausibly account for why the low variance models achieved the strongest accuracy on Context B. However, although this redundant coding strategy may help stabilize performance within a single context amidst overall environmental instability, it ultimately proved ineffective because it limited the rapid adaptability needed to operate in a dynamic environment with multiple context-dependent structures, resulting in a failure to express knowledge of both contexts.

Mirroring the diversity of these computational profiles, humans also exhibited considerable variability. Although performance on context-dependent trials was significantly above-chance at the group level, some participants exhibited little or no learning (akin to the high-weight models) while others showed stronger learning of one of the contexts (similar to the low-weight models). Just as some GRU models required more exposure to learn both sets of associations, certain individuals may also need more input to reach stable learning. A promising future direction is to identify model parameters that reflect these individual differences and predict how quickly a learner converges on context-dependent associations, potentially linking such parameters to developmental changes in learning efficiency.^59^

Taken together, our findings demonstrate that humans can spontaneously resolve conflicting, context-dependent associations from passive exposure alone – even in the absence of explicit instructions, self-pacing, feedback, or contextual cues. The finding that explicit signaling offered no advantage over entirely latent context exposure further highlights the robustness of this incidental learning mechanism, suggesting that temporal statistics alone are sufficient to drive contextual inference. Our neural network modeling provides a mechanistic account for this capacity, showing that successful adaptation relies on the emergence of distributed representations that are influenced by weight initialization parameters, which we believe are a reasonable proxy for humans’ inductive biases. This representational strategy not only maintains information from multiple contexts, even when the associative structure directly conflicts across contexts, but also quickly accommodates context changes. These findings suggest that the human brain may rely on similar mechanisms to flexibly manage latent contextual shifts and support adaptive prediction in dynamic environments.

## Materials and Methods

### Participants

Participants were recruited via the UCLA Psychology Department subject pool and completed the experiment in-person for course credit. All participants provided informed consent in accordance with protocols approved by the UCLA Institutional Review Board (IRB#22-001719). Inclusion criteria of aged between 18-40 years, native English speaker, and normal or corrected-to-normal vision with contacts (no glasses) were confirmed before commencing data collection. Our goal was to obtain useable data from 50 participants for each of the two experiments (Expt. 1: Unsignaled context; Expt. 2: Signaled context), so enough participants were collected to reach our data quality thresholds of 90% of trials responded to and 85% accuracy on trials during the learning phase. These inclusion criteria were enforced to ensure that data analyses focused on participants who were engaged during the learning phase of the experiment. Our final sample included 50 participants for Expt. 1 (33 F / 17 M; mean age = 20.5 years) and 50 different participants for Expt. 2 (40 F / 8 M / 2 Non-Binary; mean age = 20.0 years).

### Materials

The experiment was coded and run with PsychoPy version 2024.2.4^60^ on a Mac Mini. Stimuli were displayed on a DELL P2422HE monitor with 1920 by 1080 pixel resolution and screen size of 23.8 inches, which participants viewed from a fixed distance with their head stabilized with a forehead and chin rest. An EyeLink 1000 eye tracker (SR Research) captured gaze location while participants completed the experiment, but eye tracking data are not reported here. Experiment stimuli were drawn from a set of objects created using Blender 2.48.^61,62^ The stimuli were visually distinct in terms of shape and color and were novel to participants. Images were resized to be 350 pixels wide. A small “×” or “+” symbol was subtly embedded onto each object using slight color contrast such that the mark was visible but did not obstruct recognition of object shape.

### Learning phase

In the first phase of the experiment, participants were exposed to a sequence of objects presented individually. The objects were presented in four different locations on the screen with a width of 350 pixels and centered 300 pixels above, below, right, and left of the center of a gray screen. At each object presentation, the three positions not occupied by the current object were filled with phase-scrambled versions of other objects cropped into circles with diameter of 300 pixels (visualized in *SI Appendix,* Figure S2). The experimental manipulation of object location was included to enable potential analyses of spatial location-based learning as indexed by anticipatory eye movements. However, because the eye tracking data did not yield clear or interpretable effects, we focus all analyses on object identity and omit spatial position from further consideration, as well as from the task depiction in Fig. 1. Before beginning the learning phase, participants were instructed that parts of the sequence might become familiar over time and that they would later be asked questions about the objects they had seen.

Unbeknownst to the participants, the objects were organized into two sets (or contexts) of 5 pairs of objects. The same object set was used for all participants but were randomly assigned to each pair position, and each object maintained either first-of-pair (item 1) or second-of-pair (item 2) position in the pair across contexts. One pair was context-independent, meaning the same two objects were paired in both contexts. The other four pairs were context-dependent. Three of these pairs consisted of the same set of six objects across both contexts, but the second item associated with each first item was dependent on context. For example, Object X is paired with Object Y in Context A but with Object Z in Context B. The last context-dependent pair shared the same first object across both contexts but the paired second object was specific to each context (e.g., only appeared in that context and not the other). In this way, these four context-dependent pairs shared a first item across contexts but the second item of each pair was dependent on context. In total, a set of 11 unique items were used to instantiate the five pairs in each context.

Throughout this learning phase, participants were tasked with responding to whether the object onscreen was marked with an “×” or “+”. Therefore, reaction time to the perceptual question could be evaluated as online measures of pair structure learning. Objects were presented for 1200ms with a 450ms interstimulus interval.

The two experiments differed on with respect to whether context was Unsignaled (Expt. 1) or Signaled (Expt. 2) with a border around the objects that was white or black depending on the context.

### Two-alternative forced choice (2AFC) task

In the first of three test tasks immediately following this learning phase, participants completed a two-alternative forced choice (2AFC) task. Because the object associations were dependent on active context for all but the one context-independent pair, on each 2AFC trial participants were presented with a sequence of seven objects (consisting of three pairs from one of the contexts and the first item of the test pair) before being presented with two side-by-side alternatives as to which object they think should come next (one was the correct paired associate of the test pair and the other was a lure). Objects were presented with the same timing as used during the learning phase in the sequence, and participants were given unlimited time to make a choice between target and lure. Participants completed a total of 54 questions: 6 of these questions evaluated the context-independent pair, while 48 questions evaluated context-dependent associations. The 48 questions probing context-dependent associations could either feature a lure object that was the correct paired associate in the other context (direct-conflict; 16 questions), or a lure that was any other item (indirect-conflict; 32 questions). The “×” and “+” markings were removed from the objects to make clear that participants no longer were required to respond to the perceptual question. For the Unsignaled experiment, no explicit context cues were provided; for the Signaled experiment, the border around the objects was colored white or black on each trial to cue contexts. After making each 2AFC judgment, participants were prompted to rate their confidence in their decision from 1-4.

### Structure knowledge probe

After completing all 2AFC trials, participants were prompted to answer some questions about what they learned during the experiment. First, they were asked to respond yes or no to whether they observed any predictable patterns in the experiment. Second, they were asked to describe any patterns they observed in the sequence. Third, they were asked to describe any rules that governed which object would come next in the sequence. The idea was to progressively prompt participants to indicate any knowledge of the pair structure underlying the sequence they observed that were increasingly straightforward to get an idea of how much knowledge was explicit.

### Pair reconstruction task

The last task allowed participants to demonstrate explicit knowledge of pairs. Participants were presented with a bank of all 11 objects at the top of the screen and provided with 20 sets of two empty squares side-by-side presented in 4 rows of 5 columns. Participants were instructed to organize the objects into related pairs by placing one object in each square of a pair, with the left and right positions corresponding to the first and second items in the pair. Participants were told that each item could be used more than once and that they did not have to fill out all of the pairs.

### Data cleaning

Data exclusion criteria were enforced to ensure that participants were engaging with the experiment during the learning phase. As such, two criteria were enforced: response rate of more than 90% and accuracy of more than 85% on all responses throughout the learning phase. Data collection continued until 50 useable participants were collected for each experiment.

### Neural network architecture

Recurrent neural network models with gated recurrent units were implemented with PyTorch v2.0.1 ^63^. Such models have previously been used to explore context-dependent associative learning from sequences ^22,27^. Each model had the same architecture: an 11-node input layer with dimensionality of 11 (equal to the number of objects included in the study), a hidden layer of 150 nodes (GRU performance with different hidden layer sizes presented in *SI Appendix*, section S5), and an 11-D output layer again to match dimensionality of one-hot object vectors. Learning rate was held constant at 0.001, and model weights were updated after each training sample using the Adam algorithm of gradient descent and cross entropy loss. Default parameters were used unless otherwise noted.

### Training

A unique 1600-object sequence was generated for each model in the same way as for human participants. Each neural network received one object at a time and was trained to predict the identity of the next object in the sequence. Although the sequence was constructed using embedded object pairs, models received no information about this underlying structure. That is, the model made predictions at every time step (1599 samples for the 1600-object sequence) and had no awareness of pair boundaries. The same sequence was used for all epochs of training, with the hidden state was reset at the start of each epoch and between blocks (every 400 samples) in recurrent models to emulate the breaks taken by human participants.

### 2AFC task

After each epoch of learning, model weights were frozen, and the models were evaluated using a 2AFC test designed to mirror the testing procedure of the human participants. A unique set of 2AFC test questions was generated for each model in the same way as for human participants. Before each trial, hidden layer activity was reset to zero. Then, a sequence of three pairs from one of the contexts was presented as the hidden state evolved, allowing the model to infer the active context based on the sequence. Finally, the first object of the test pair was inputted, and the model’s prediction of the ensuing item was evaluated. Accuracy was determined by whether the probability assigned to the correct paired associate was higher than that to the lure. In most analyses, accuracy is evaluated separately for the context-independent, indirect-conflict context-dependent, and direct-conflict context-dependent question sets to capture how well the models handle conflicting information across contexts and maintain knowledge of stable, context-independent relationships.

### Single epoch analyses

We tested the GRU’s ability to learn the task as the variance of the uniform distribution used to initialize the hidden layer’s weights was increased. The uniform distribution was centered at zero with positive and negative bounds of 0.08 (default for PyTorch with 150 nodes), 0.2, 0.4, 0.6, 0.8, 1.0, 1.2, and 1.4. Fifty independent GRU models with different weight initialization randomizations were trained and tested, and learning measures across these models were averaged to ensure robust performance estimates of each weight initialization category.

### Learning trajectory analysis: Context switch latency

To assess how quickly the neural network models adapted to a context change, we developed a switch latency measure. We devised a stringent operationalization of switch latency as the number of first-of-pair (item 1) items (e.g., the item whose model output captures the within-pair transition prediction) a model processed after a context switch before achieving perfect accuracy on all remaining item 1 samples in that 50-pair context exposure. Because only the model outputs of item 1 training samples are predictable and thus learnable, they served as a measure of adaptation to a new context. Switch latency was calculated for all 16 context exposures (8 per context) and averaged within each block (4 context exposures), yielding a single switch latency value per block. We averaged this measure across all 50 trained models for each weight initialization condition.

### Hidden layer analyses: Context representation strategies

To understand how context information was represented across the hidden layer, we quantified two complementary properties of the activations: the extent to which context sensitivity was localized to a small subset of units (akin to individual “context cell” neurons that code for which context is currently active), and the degree to which the currently active context was expressed a as distinctive pattern of activity across many units. These analyses were conceptually motivated by prior work distinguishing sparse and distributed coding strategies in hippocampal and connectionist models.^35,41^ Our goal was to determine whether these representational properties could explain 2AFC task performance across all individual GRU model instances of the weight variance configurations.

A sparse representation describes when context sensitivity is confined to a relatively small subset of hidden layer units, while the vast majority remain inactive or insensitive. To investigate sparse context representations in the GRU’s hidden layer, we first used a one-way ANOVA to estimate the difference in activation when processing inputs from Context A and Context B during the final quarter of training (block 4) for each of the 150 hidden layer nodes. We then counted the number of nodes that showed a significant activation difference. We applied a Bonferroni correction within analysis of each model to control for Type I errors of the 150 comparisons were performed. The corrected significance threshold was computed by dividing the original alpha level (0.05) by the number of comparisons (150), yielding an adjusted significance level of p < 0.00033. Based on this threshold, we determined that F_crit_(1,398) = 12.75 and calculated the sparse representation index as the proportion of hidden layer nodes that did not show a significant difference in activation between contexts, such that a larger value reflects a sparser context representation.

A distributed representation was computed using a representational similarity analysis (RSA; ^64^) focused on the hidden layer activations after processing the first item of each pair (capturing the context-dependent prediction) during the final quarter of training (block 4). This included a total of 200 hidden state samples (100 per context). These activations were divided into two split-halves, each containing 10 samples for each of the five pairs per context. These samples in each split-half were evenly divided into those drawn from the first half of a context exposure and those from the second half, controlling for any strengthening of context representation over time. We averaged the hidden state activation within each node for each object within each context. We then computed the Pearson correlation coefficient for all pairwise comparisons of objects within and across contexts, producing an RSA matrix. This matrix compared the split-half object representations, with one half plotted along the x-axis and the other along the y-axis, and there were 10 cells along each axis for each of the five pairs viewed in each context. The upper-left and lower-right quadrants contained correlations between the five pairs from the same context. The upper-right and lower-left quadrants contained correlations between the same objects when viewed in opposing contexts. To quantify the distributed representation, we calculated the difference between the average within-context and between-context correlations for each object, normalized by subtracting the average within-context correlation from one. This normalization penalized models with lower within-context stability because a larger denominator as a result of lower within-context similarity would decrease the overall distributed representation index, ensuring that observed differences in between-context representation were not artifacts of noisy or unstable object representations.

### Hidden layer analyses: Lesion analysis

To further understand how the GRU model’s hidden layer representations supported successful context-dependent learning, we conducted an intervention analysis. For all weight configurations, we trained a new set of 50 models with the same single-epoch training procedure and then systematically tested their performance on the 2AFC task while “lesioning” (zeroing out) subsets of hidden layer nodes. Importantly, these nodes were active during training on the 1600-object sequence, and the lesioning intervention was applied only immediately prior to the 2AFC testing phase.

We first calculated the absolute context sensitivity of each hidden layer node using a one-way ANOVA of activations between Context A and Context B during the final quarter of training (block 4), in the same way as computing the sparse representation index. Nodes were then ranked from the most context-sensitive (largest *F*-statistic) to least sensitive. During the 2AFC task, subsets of hidden layer nodes were progressively lesioned, beginning with the most context-sensitive nodes. We evaluated models under the following lesioning conditions: 0 (no nodes lesioned to obtain baseline performance estimate), 1, 5, 10, 25, 50, 75, 100, 125, 130, 135, 140, 145, and 150 (all nodes lesioned with expectation of chance performance). We report the average 2AFC performance on Context A, Context B, and context-independent question sets, expressed as both obtained accuracy and the change in performance relative to the no-lesion baseline (e.g., when no intervention is applied).

### Statistical Analysis

#### 2AFC task

2AFC task performance was assessed by evaluating accuracy and average confidence rating on subsets of 2AFC questions. Group-level accuracy was tested against 50% chance using a one-sample Student’s *t*-test with Holm-Bonferroni correction applied for three comparisons (Context A, Context B, and context-independent trials) within each experiment.

We conducted a mixed-design ANOVA to examine the effects of experiment (between-subjects factor: Expt. 1 versus Expt. 2) and context-dependence (within-subject factor: context-dependent versus context-independent trials) on 2AFC accuracy using the Python pingouin package. To assess whether context-dependent 2AFC accuracy was statistically equivalent between experiments, we used Bayesian estimation with a region of practical equivalence (ROPE) approach. We computed the posterior distribution of the mean difference in accuracy between Expt. 1 and Expt. 2 and quantified the proportion of the posterior mass falling within a predefined ROPE of [-5%, 5%]. This ROPE was selected to reflect the smallest effect size of interest, consistent with typical variability in task accuracy in statistical learning literature. Posterior distributions were estimated using the PyMC package.

#### Online learning assessment

We quantified online learning using participants’ RTs during the learning phase, in which they indicated whether each object contained an “×” or “+”. For each block, we computed an anticipation score as the average RT to the second item of each pair subtracted from the average RT to the first item of each pair. This metric captured facilitation for predictable second items while controlling for overall RT drift throughout the session, as first items follow unpredictable transitions. Positive values indicate faster responses to second items relative to first items.

To assess changes in online learning across blocks, we applied linear contrast with weights [-3, - 1, 1, 3] to the blockwise anticipation scores for each participant and tested the group-level difference from zero using a two-tailed one-sample *t*-test for each experiment.

#### Multivariate regression analysis of representation strategies

The two measures of hidden layer activity – sparse representation index (proportion of hidden layer nodes that do not show significant activation difference by context) and distributed representation index (within-versus between-context correlation differences) – were used as predictors in multivariate regression models aimed to explain the variance of 2AFC context-dependent accuracy, context-independent accuracy, and accuracy difference between contexts. These indices were first computed for each of the 50 models instantiated with each of the eight weight initialization configurations, and then were averaged within each configuration. The resulting eight values for each predictor were z-scored to enable comparison of beta coefficients across predictors. By including both metrics in the same regression models, we assessed their unique contributions to task performance. This allowed us to evaluate whether sparse or distributed representations were more predictive of learning outcomes, providing insight into the mechanisms underlying the GRU model’s ability to process and adapt to context-dependent associations.

## Acknowledgements

F.P. was supported by the National Science Foundation Graduate Research Fellowship Program under Grant Nos. DGE-2034835 and DGE-2444110.

## Author Contributions

Conceptualization: FCP, JR, HL; Methodology: FCP, JR, HL; Software: FCP; Formal analysis: FCP; Visualization: FCP; Supervision: JR, HL; Writing – original draft: FCP; Writing – review & editing: FCP, HL, JR.

## Supplementary Information Appendix

**Fig. S1.**
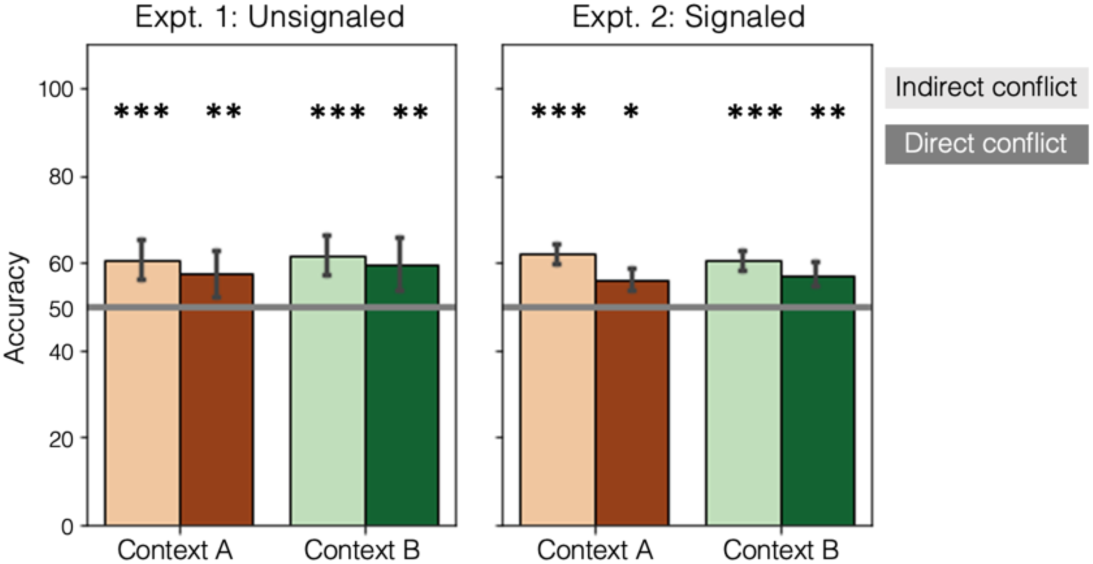
2AFC test performance by direct and indirect conflict question subsets. Bar height reflects group average on 2AFC context-dependent question subsetted by indirect (light coloring) and direct (dark coloring) conflict for Context A (left, orange bars) and Context B (right, green bars). ***p<0.001; **p<0.01; *p<0.05.

**Fig. S2.**
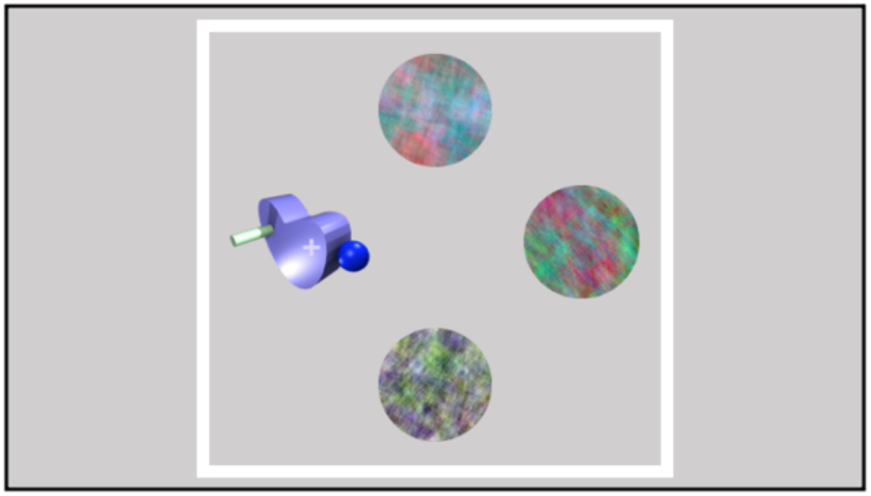
Visualization of stimulus presentation during learning phase. Each trial of the learning phase featured four stimuli arranged as depicted, with one object of interest (on which participants needed to make an × /+ judgment) and three circular phase-scrambled objects presented in the remaining positions. A black or white border was present during Expt. 2. Visualization of objects and border is to scale.

**Table S1.** Reaction times (mean ± standard deviation) in milliseconds by block for context-dependent pair objects. Item 1 is the first, unpredictable element of each pair; Item 2 is the second, predictable element informed by the associative expectation.

|  | Expt 1: Unsignaled |  | Expt 2: Signaled |  |
| --- | --- | --- | --- | --- |
|  | Item 1 | Item 2 | Item 1 | Item 2 |
| Block 1 | 712 ± 72 | 727 ± 75 | 710 ± 67 | 713 ± 57 |
| Block 2 | 668 ± 77 | 679 ± 84 | 662 ± 76 | 664 ± 64 |
| Block 3 | 650 ± 76 | 655 ± 85 | 648 ± 77 | 643 ± 62 |
| Block 4 | 639 ± 74 | 639 ± 83 | 631 ± 71 | 626 ± 62 |

### S1 Confidence judgments and explicit knowledge assessments

We examined whether participants’ confidence ratings on the 2AFC task were related to their accuracy. Confidence was significantly higher for accurate than inaccurate 2AFC responses for context-dependent trials in both experiments (Unsignaled: *t*(49) = 4.1, *p* < 0.001; Signaled: *t*(49) = 3.96, *p* < 0.001) and on context-independent trials in Expt. 1 (Unsignaled; *t*(37) = 2.6, *p* = 0.013) but not Expt. 2 (*t*(39) = 0.38, *p* = 0.7) (Fig. S3A). Participants who responded entirely correctly or incorrectly were excluded from analysis; all participants had mixed accuracy on context-dependent trials. Overall, mean confidence ratings were around or below the midpoint of the scale, indicating generally low subjective certainty during the 2AFC task.

Following the 2AFC task, participants completed two additional assessments designed to measure explicit knowledge of the temporal structure: a Structure Knowledge Probe and Pair Reconstruction Task.

For the Structure Knowledge Probe, binary performance was assessed by manually evaluating whether participants articulated explicit awareness of temporal pair structure in their written responses. This measure did not evaluate knowledge of the dual context structure (e.g., participants did not need to articulate awareness that there were two distinct contexts where the associative pairings changed). Two independent raters coded all responses with 91% agreement; discrepancies were resolved by deferring to the more senior grader. Explicit knowledge of the pair structure was identified in 34.0% of participants in Expt. 1 and 40.0% in Expt. 2.

Performance on the Pair Reconstruction Task varied because participants could report between 1 and 20 pairs. To estimate chance performance, we implemented Monte Carlo simulations where 1,000 simulations were run for each possible number of reported pairs (k = 1-20). In each simulation, an object was sampled from the 11 unique objects with replacement between pairs but without replacement within a pair (e.g., no pair comprised of the same object). This produced a null distribution of proportion correct entries for each k expected by chance. This empirical approach matches the analytical solution: there were 9 correct pairs (because one pair was context-independent and thus correct in both contexts), and the probability of guessing one correct pair by chance was 1/110 (choosing 2 of the 11 objects without replacement). Thus, the probability of guessing one of the 9 correct pairs was 9/110, or 8.2%.

Participants reported an average of 7.4 ± 4 in Expt. 1 and 7.9 ± 4 in Expt. 2 (Fig. S3B). Group-level significance was calculated as the average number of correct context-independent pairs was greater than expected by chance over the simulations of all possible pair entry counts.

Context-dependent pair entry performance was non-significant for both experiments (Unsignaled: mean = 2.06 pairs; p = 0.11; Signaled: mean = 2.08 pairs; p = 0.11; Fig. S3B). Moreover, only 36% of Expt. 1 participants and 34.7% of Expt. 2 participants (e.g., 17 out of 49 participants; one participant did not complete this portion of the experiment) reported the context-independent pair.

Taken together, these results indicate that most participants had little to no explicit knowledge of the temporal pair structure: they were generally unable to recall the context-independent pair, articulate the underlying pair structure, or reconstruct the context-dependent associations. Thus, the significant 2AFC performance reflecting context-dependent learning is unlikely to have been driven by explicit awareness.

**Fig. S3.**
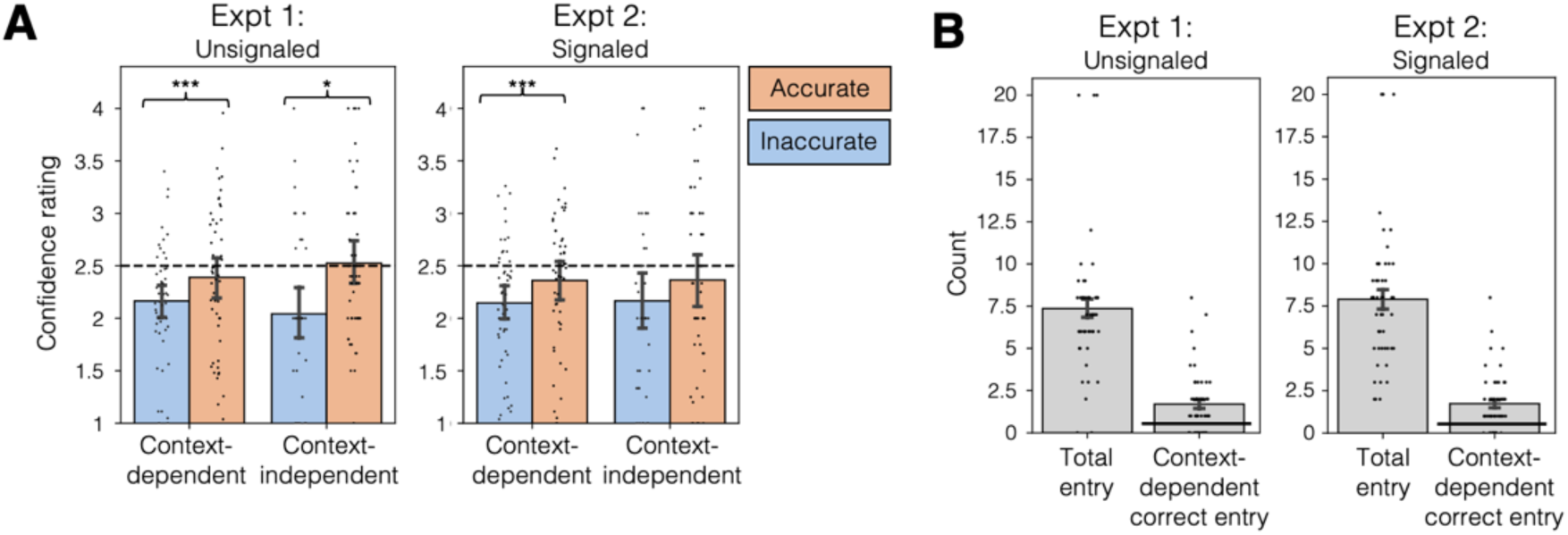
Metacognitive awareness assessment results. *(A)* Average 2AFC confidence rating for context-dependent (left) and context-dependent (right) questions for accurate (pink) and inaccurate (blue) question responses. Horizontal dashed line indicates midpoint of the confidence scale, and error bars reflect SEM. *(B)* Pair Reconstruction Task performance: bar height reflects group average number of total pairs reported (left) and correct context-independent pairs reported (right). Horizontal line reflects chance performance based on Monte Carlo simulations; error bars reflect SEM.

### S2 Model architecture comparison

In the main paper, we analyze a GRU model with training that was constrained to a single epoch, equivalent to the total sequence exposure of each human participant. Here, we justify that decision with comparison to two simpler models: a feedforward neural network (FFNN) and a vanilla recurrent neural network (RNN) that lacked gated recurrent units. One learning phase sequence and one set of 2AFC questions were generated for each model in the same way as for human participants (1,600 objects), and each epoch of training consisted of updating model weights to predict the next item in this sequence, followed by an assessment of 2AFC accuracy with frozen weights. For each model, we continued this process for a total of 50 epochs (i.e. 50 times the sequence exposure given to human participants). All models here used the default PyTorch weight initialization where weights are drawn from a uniform distribution bounded by plus or minus the inverse of the square root of the layer size, which was 0.08 for the hidden layer.

The simplest architecture, the FFNN, achieved an overall context-dependent accuracy of almost 75% (Fig. S4A). However, this performance was entirely driven by near-perfect accuracy on Context B, the most recently trained context, while accuracy on Context A remained near chance. This indicates that the FFNN retained knowledge only about the most recent associations, completely overwriting previously learned, conflicting ones—a hallmark of catastrophic interference.

RNNs improve on feedforward model capabilities by incorporating information from past hidden states with the current state, enabling them to process sequential input. However, the RNN showed no improvement in overall context-dependent accuracy compared to the FFNN (Fig. S4B). While Context A performance did increase over learning, this improvement came at the expense of Context B performance, suggesting the RNN is also prone to interference.

The RNN with GRUs, an advanced RNN variant, overcomes the limitations by using update and reset gates to manage long-term dependencies more effectively. Initially, GRU performance was comparable to the FFNN (Fig. S4C). However, with extended training (approximately 20 epochs; i.e., 20 times the exposure of human participants), the GRU achieved comparably high accuracy on both Context A and Context B. This performance likely stems from the GRU’s architectural advantages. The update gate controls how much new input influences retained memory, allowing the model to ignore unreliable input (such as noisy between-pair transitions). The reset gate allows the selective clearing of irrelevant information in response to context changes, thereby avoiding interference from outdated associations. Given that the GRU model provides the best account for context-dependent learning in humans, we used the GRU model for all of the single-epoch modeling analyses reported in the main text.

**Fig. S4.**
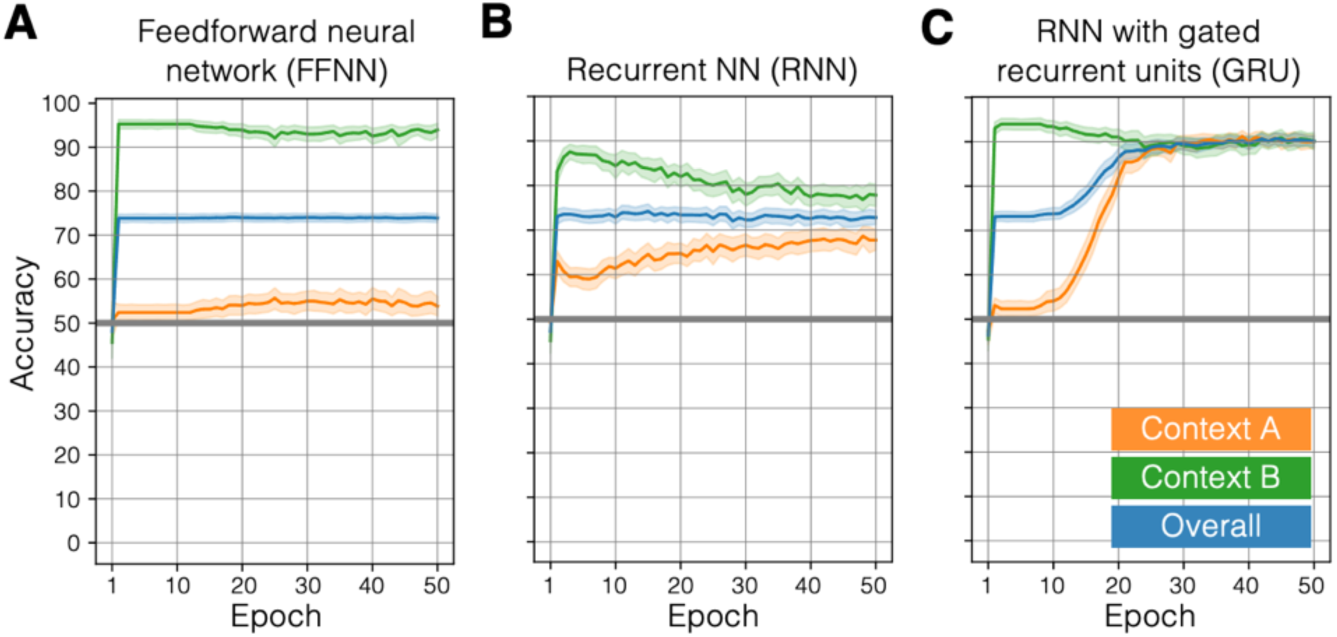
2AFC task performance for three neural network model classes. *(A-C)* 2AFC performance accuracy on context-dependent questions averaged for 50 individual models for each architecture after each of 50 epochs of training; accuracy is plotted separately for Context A (orange), Context B (green), and both contexts combined (blue) for each model class: *(A)* simple feedforward neural network (FFNN), *(B)* vanilla recurrent neural network (RNN), and *(C)* recurrent neural network with gated recurrent units (GRU). All models achieved perfect accuracy on context-independent questions after the first epoch (not pictured).

### S3 GRU performance with perceptual object representations

The main paper used one-hot vector representations for each object in the modeling analysis. This choice ensured that all objects were represented equally and orthogonally, such that any structure emerging in the hidden layer reflected purely learned associations rather than preexisting similarities among the inputs. Here, we present the same analysis using perceptual object representations that more closely approximate the visual experience of human participants in the task. Perceptual object representations were generated by inputting each object image (without the overlaid plus or minus symbol) into AlexNet (1) and then applying PCA to reduce the dimensionality to 11 dimensions, matching the number of input and output dimensions of the original model. GRU networks were trained on the same context-dependent sequential prediction task as in the main text, using cosine similarity as the loss function, and assignment of objects to specific pairs was randomized for each model in the same way as for human participants.

As shown in Fig. S5, the relationship between initialized weight variance and 2AFC accuracy retained the same non-monotonic profile observed in the models that used one-hot input coding (Fig. 4A). In addition, 2AFC performance was higher for all question subsets. This improvement is unsurprising: the use of AlexNet embeddings introduces a strong visual prior that allows the model to exploit shared perceptual features when making predictions, thereby obscuring the interpretability of how the hidden layer activity represents the task’s temporal associative structure. For example, the model could leverage arbitrary similarities in dimensions such as shape and color to bias its 2AFC responses. In contrast, one-hot encodings constrain all non-active dimensions to zero, ensuring that any hidden layer structure arises exclusively from learning the task’s associative regularities. Taken together, these results confirm that the core finding of optimal task performance emerging at moderate weight initialization variance holds regardless of input representation. We therefore focus analysis on models trained with one-hot object encodings because they provide a controlled representational space in which hidden layer structure reflects learning task’s associative structure rather than preexisting perceptual relationships among stimuli.

**Fig. S5.**
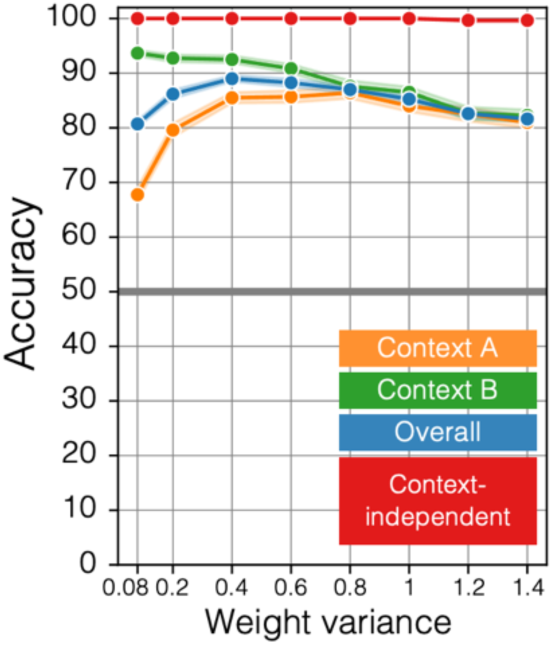
2AFC performance with perceptual object embeddings. 2AFC accuracy (y-axis) on context-dependent test trials for GRU models with weights initialized with increasing variance along the x-axis color-coded by question category.

### S4 GRU performance with a fully overlapping stimulus set (no context-specific objects)

The pair assignment across contexts for both models and human participants included one possible caveat to our claim of latent context-dependent learning: for one of the context-dependent pair types, the second (paired) item appeared in only one of the contexts while the first item occurred in both (visualized in bottom row of Fig. 1B). This structure still required humans and the models to update their prediction of the second item based on the inferred context (consistent with all other context-dependent pairs), but also meant that the context-specific second item could have been used as a context cue independent of recent sequence history (Expt. 1) and/or border color (Expt. 2). In other words, an encounter with a context-specific object could be an indicator that the state of the world has changed and thus that one’s associative predictions should be updated. We note that our decision to include this pair type was motivated by our intention to collect fMRI data with this paradigm, which will allow us to assess changes in neural representational geometry when an object’s associative identity remains constant across contexts, providing a baseline for evaluating relative changes in other pair conditions.

To evaluate whether the presence of context-specific objects influenced model learning, we ran neural network simulations in which such pairs were removed and replaced with context-dependent pairs for which both objects could occur in either context. These models were trained no the same task, but the context-dependent pairs were reconfigured to maintain the overall object set, with second-item assignments shuffled across contexts. Model input and output dimensions were therefore reduced to 10, corresponding to the 10 unique object encodings needed to instantiate this modified pair set. All other training parameters and analysis of 2AFC performance were identical to those in the main text.

As shown in Fig. S6, model performance across weight initializations closely mirrored the results of the main analysis (Fig. 4A), indicating that learning dynamics and context-dependent accuracy were unaffected by the presence or absence of the context-specific object. This indicates that such objects did not serve as reliable context cues for our models.

**Fig. S6.**
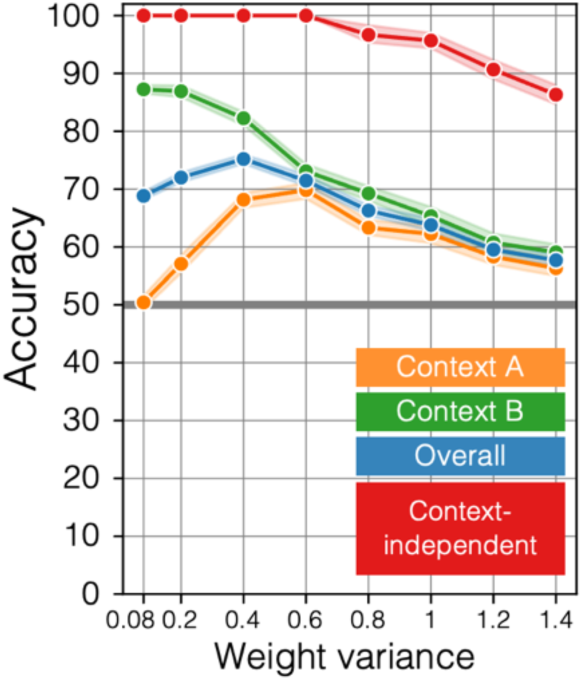
2AFC performance with no context-specific objects measured by weight variance. 2AFC accuracy (y-axis) on context-dependent test trials for GRU models with weights initialized with increasing variance along the x-axis color-coded by question category.

### S5 Determination of hidden layer size

The main paper analyzes a GRU model with 150 hidden layer units. To assess whether model capacity influenced learning performance, we trained GRU models with reduced hidden layer sizes of 50 and 100 units. As shown in Fig. S7, all models ultimately achieved comparable performance converging to approximately 90% accuracy. However, models with fewer hidden layer units exhibited slower learning trajectories, requiring more training to reach the same level of performance as the model with 150 units. These results suggest that while increasing the number of hidden units accelerates learning, overall task performance is largely independent of model size.

**Fig. S7.**
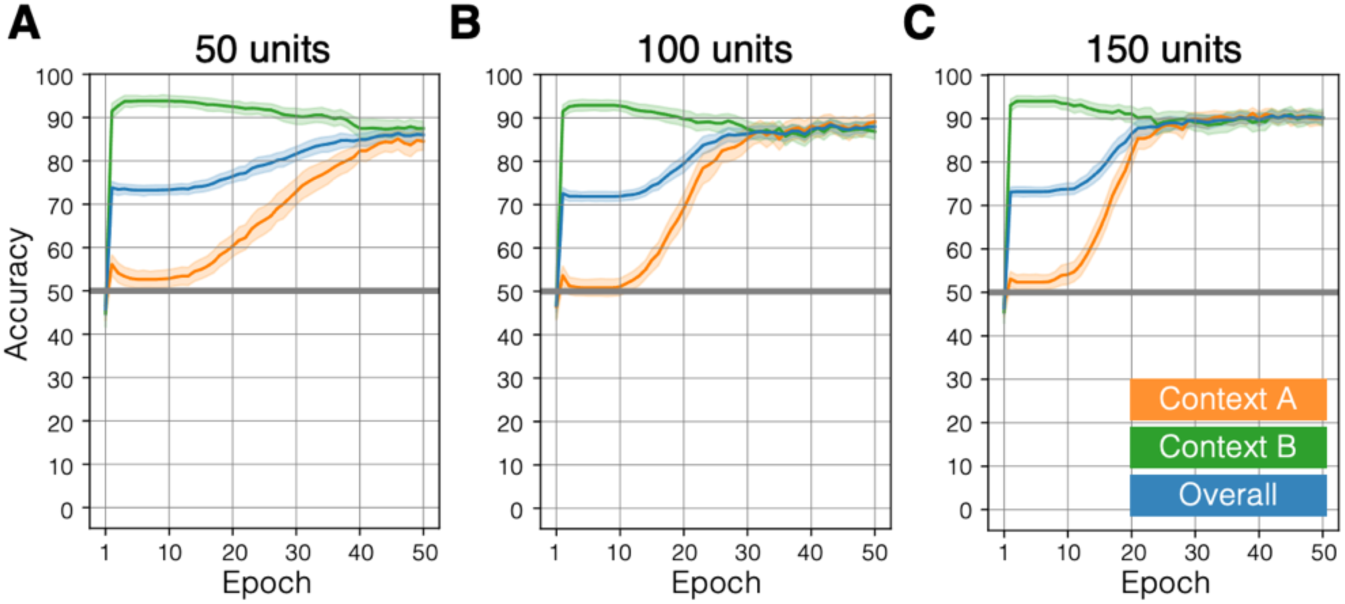
2AFC task performance for GRU models with varying hidden layer sizes. *(A-C)* 2AFC performance accuracy on context-dependent questions averaged for 50 individual models for each architecture after each epoch of training for Context A (orange), Context B (green), and both contexts combined (blue) for GRU models with *(A)* 50 hidden layer units, *(B)* 100 hidden layer units, and *(C)* 150 hidden layer units.

